# Mitigating Permittivity–Charge Cross-Sensitivity in Biomolecular Sensing: Electrostatic Decoupling and Feedforward Biasing in an Asymmetric Dual-Gate a-IGZO FET

**DOI:** 10.64898/2026.10.15.759373

**Authors:** Bakhsh Mohammed Mahanna, Muntasir Amin, Muhammed Shafayat Jamil, Mohammad Masum Billah

## Abstract

Permittivity-based biomolecular identification is inherently ambiguous in real-world sensing environments because analyte charge also influences the measured electrical response. Different combinations of charge and permittivity can produce similar electrical responses, so a change in the measured response cannot be uniquely attributed to permittivity. This work combines electrostatic decoupling through device engineering with charge-conditioned feedforward biasing to mitigate this cross-sensitivity and recover both analyte properties through direct drain-current readings from a single transistor. We provide a proof of concept through technology computer-aided design (TCAD) simulations of an asymmetric dual-gate, dual-nanogap amorphous indium– gallium–zinc oxide (a-IGZO) transistor. Gate-insulator dielectric engineering and nanogap optimization establish complementary drain-current responses, with the bottom gate predominantly sensing analyte charge and the top gate predominantly sensing permittivity. Feedforward biasing further suppresses residual charge interference in the top-gate response. First, the bottom-gate drain current provides an estimate of analyte charge. This estimate and a preliminary top-gate current reading are used to select the voltage for the final top-gate measurement. The drain current measured at that voltage then determines permittivity. Three sequential current readings thus provide permittivity-based biomolecular identification without prior knowledge of analyte charge, alongside complementary charge measurement. Mean charge and permittivity extraction errors remain below 1.8% and 0.6%, respectively, across the calibration and independent test datasets.

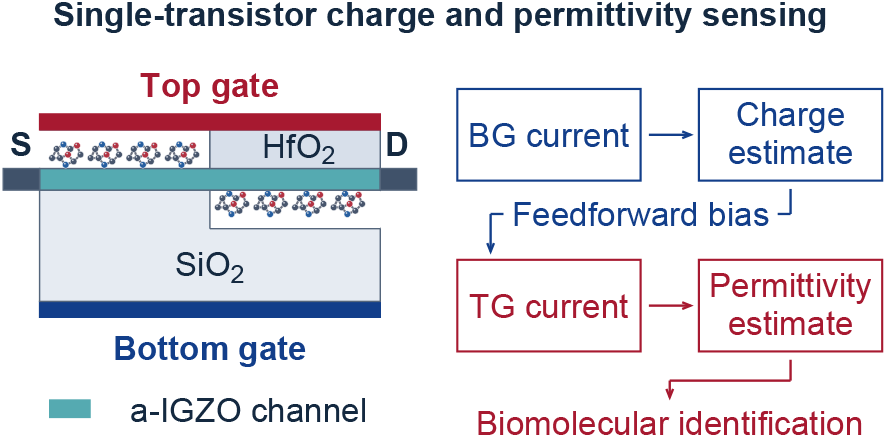

## I. Introduction

Dielectric-modulated field-effect transistors (FETs) detect biomolecules through changes in their electrical response, and experimental nanogap devices and separated double-gate platforms have established dielectric-sensitive detection through electrical responses [1], [2]. However, the measured current, threshold voltage, or transconductance can respond to both biomolecular permittivity and charge. A change attributed to permittivity may therefore originate partly from charge, creating ambiguity when both properties are unknown. This charge– permittivity interference presents a central challenge to reliable permittivity-based biomolecular identification and motivates the independent measurement of both properties.

Assuming charge neutrality can eliminate this ambiguity under controlled conditions, but such an assumption does not necessarily hold in practical sensing environments. Biomolecular charge depends on ionization, pH, and screening [3], [4]. Different charge–permittivity combinations can produce similar responses. For example, a neutral analyte with higher permittivity may produce a response resembling that of a charged analyte with lower permittivity when the charge contribution compensates for the dielectric difference. Consequently, a permittivity-sensitive output alone cannot reliably distinguish these possibilities without additional information about charge. Addressing this challenge requires independent determination of both analyte properties rather than merely improving sensitivity to either one.

Previous studies establish charge–permittivity interference. At a fixed dielectric constant of *k* = 5, Narang *et al*. reported that a charge density of −10^12^ cm^*−*2^ reduces ON-current by approximately 50% in a conventional dielectric-modulated metal–oxide–semiconductor FET (DM-MOSFET) and by an order of magnitude in a dielectric-modulated tunnel FET (DM-TFET) [5]. Their subsequent study showed that charge contributions can dominate dielectric modulation at lower permittivities [6]. Singh *et al*. studied single-gate-operated sensors and found that their back-gate structure is sensitive to negatively charged biomolecules, limiting neutral-analyte detection, whereas their front-gate structures respond to both permittivity and charge [7]. Their study, however, compares sensitivity rather than extracting both simultaneously varying properties. Similarly, Narang’s sensitivity studies, Srikanya’s amorphous indium–gallium–zinc oxide (a-IGZO) charge and dielectric analysis, and gate-stack and nanogap-layout studies characterize mixed charge–permittivity responses but do not demonstrate mitigation of this cross-sensitivity or recovery of both unknown analyte properties, and assume controlled charge environments or neutral analytes [6], [8]–[10]. These studies establish the significance of charge–permittivity interference but leave the challenge of independently determining both properties largely unaddressed.

Because the effective analyte charge varies with the sensing environment, accurate permittivity-based identification under realistic conditions requires the charge contribution to be measured or accounted for alongside the dielectric response [3], [4], [6]. Kim *et al*. established a precedent for joint extraction using an asymmetric Si-nanowire double-gate transistor [11]. Their method extracts permittivity through simultaneous gate sweeps, subthreshold-slope analysis, and capacitance reconstruction using biomolecular thickness, followed by charge determination from the threshold-voltage shift. Although this work demonstrates the feasibility of jointly extracting charge and permittivity, its measurement and extraction procedure involves multiple gate-bias conditions and the determination of several intermediate electrical parameters. This highlights an important opportunity for a more direct approach that retains the ability to determine both unknown analyte properties while simplifying the measurement procedure.

We use an asymmetric dual-gate, dual-nanogap a-IGZO transistor engineered for complementary charge- and permittivity-sensitive current responses. Unlike Kim’s parameter-based extraction procedure, the proposed architecture uses dielectric-layer and nanogap engineering to mitigate cross-sensitivity at the device level before readout correction. The bottom-gate (BG) response provides charge estimation and the top-gate (TG) response provides permittivity determination through two reference current curves, while the estimated charge selects an adjusted top-gate voltage to further reduce residual interference. The resulting strategy combines electrostatic decoupling through device engineering with feedforward selection of the top-gate readout bias. The two extracted properties, (*k, N*_bio_), describe the analyte’s dielectric environment and effective interfacial charge, reflecting its polarization and electrostatic behavior, respectively [12], [13].

Together with this device-level separation of sensing responses, our approach substantially simplifies measurement and extraction relative to Kim’s. Kim obtains permittivity by determining subthreshold slopes at multiple simultaneous gate-bias ratios, locating an optimum ratio, and reconstructing the sensing capacitance using known biomolecular thickness; charge determination additionally requires a threshold-voltage-shift calculation [11]. After calibration, our method instead extracts both properties through three sequential drain-current readings. The bottom-gate current provides an initial charge estimate, a preliminary top-gate current reading determines an adjusted measurement voltage using that estimate, and the final top-gate current enables permittivity extraction. This eliminates the need for subthreshold-slope or threshold-voltage extraction, optimum gate-ratio searches, and capacitance reconstruction during sample measurement. Dielectric and nanogap engineering with calibrated voltage correction thus enables accurate permittivity extraction despite unknown charge. This Technology Computer-Aided Design (TCAD)-based proof of concept includes calibration-grid characterization and independent tests of unseen charge–permittivity combinations.

## II. Device Structure and Sensing Principle

### A. Device Geometry

One 20-nm a-IGZO channel has common source/drain (S/D) terminals, *L* = *W* = 1 *µ*m and 0.15-*µ*m contact extensions. The source-side top nanogap and adjacent HfO_2_ are each 20 nm thick. The drain-side bottom nanogap is 20 nm above a 480-nm SiO_2_ floor; both nanogaps cover half the channel (Fig. 1).

**Fig 1.**
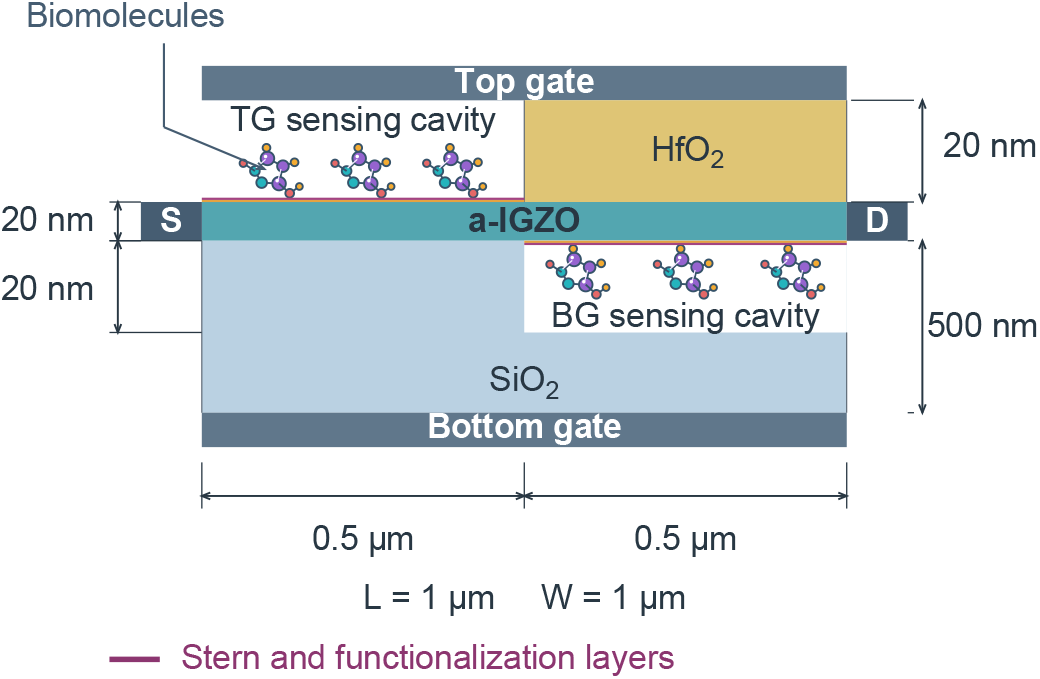
Dual-gate IGZO sensor with matched sensing nanogaps and interfacial layers; thicknesses are not to scale.

Functionalization and Stern layers of negligible thicknesses are expected in a practical device [8], [11] but are not included in the present TCAD model to simplify the proof-of-concept characterization.

### B. Nanogap-height selection and matched sensing regions

Both sensing nanogaps are 20 nm high and occupy equal 0.5-*µ*m channel lengths. This height lies within reported nanogap dimensions: 6 nm in gate-stack double-gate metal–oxide–semiconductor FET (DG-MOSFET) biosensors [14], 9 nm in double-gate junctionless FET (DG-JLFET) nanogap architectures [10], 10 nm in dual-gate transition-metal dichalcogenide FET (TMD-FET) biosensors [15], and 15 nm in dielectric-modulated impact-ionization metal–oxide– semiconductor (DIMOS) transistor biosensors [16]. Srikanya *et al*. used a 15-nm nominal nanogap and investigated thicknesses from 5 to 25 nm [8]. Kim *et al*. formed their experimental sensing gap using a 20-nm sacrificial oxide and demonstrated sequential 3-aminopropyltriethoxysilane (APTES) functionalization, biotin attachment, and streptavidin binding [11].

### C. Representation of Biomolecular Permittivity and Charge

Representing a uniformly filled sensing nanogap by one homogeneous dielectric constant is a standard idealized approach in dielectric-modulated FET modeling, used in both double-gate and IGZO biosensor studies [8], [9], [15]. In this representation: *k* specifies the nanogap dielectric response, while *N*_bio_ specifies the added equivalent sheet-charge density at the nanogap–channel interface. This charge treatment is also literature-grounded; Chowdhury *et al*., for example, represent charged biomolecules by an interface fixed-charge density [9]. It captures channel electrostatics without volumetric dielectric charge. The added *N*_bio_ is uniform along each 0.5-*µ*m sensing interface; solid-dielectric interfaces retain their original charge. Both nanogaps receive the same *k* and added density *N*_bio_; *q*_*f*_ denotes the total fixed sheet-charge density entered in TCAD, including the calibrated baseline.

At their respective channel interfaces,

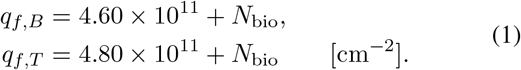

while solid-dielectric interfaces retain their baselines. The bottom and top baseline sheet-charge densities, 4.60 × 10^11^ and 4.80 × 10^11^ cm^*−*2^, respectively, were obtained by calibrating the TCAD model against the experimental transfer characteristics of Zhang *et al*.’s asymmetric dual-gate oxide thin-film transistor (TFT) [17]. The analyte-induced *N*_bio_ is added to these calibrated baseline values. Here *q*_*f*_ and *N*_bio_ are signed number densities in cm^*−*2^; their physical charge densities are *q*_*e*_*q*_*f*_ and *q*_*e*_*N*_bio_, respectively, where *q*_*e*_ is the elementary charge.

### D. Why the Two Gates Prefer Different Properties

Our device is engineered for complementary charge and permittivity responses. The unequal dielectric stacks, half-channel nanogaps, and high-*k* HfO_2_ top insulator jointly improve the overlap of top-gate current–permittivity responses across charge states. This physics-based dielectric-layer and nanogap design provides cross-sensitivity mitigation before the readout correction. The asymmetric gate stacks and nanogap layout promote electrostatic decoupling by giving the two gate currents complementary sensitivities to charge and permittivity.

Approximate local capacitances are

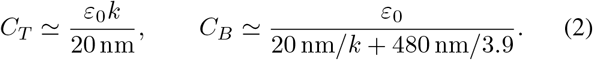

In the bottom-gate branch, the sensing gap is only 20 nm of the 500-nm dielectric thickness, with 480 nm of SiO_2_ beneath it. The SiO_2_ floor therefore dominates the series capacitance, so changing the gap’s *k* has a relatively small effect on gate coupling. Biomolecular charge, however, is placed directly at the gap–channel interface over half the channel length. Its effect on channel potential remains strong despite reduced dielectric response. The bottom-gate current consequently provides the charge-sensitive reading.

Over the top sensing region, the gap occupies the entire 20-nm gate-insulator thickness. Without a thick series SiO_2_ layer diluting its dielectric contribution, changing *k* produces a much stronger change in top-gate coupling. The top-gate current therefore provides the permittivity-sensitive reading. The shared channel still allows some charge influence on this current; the voltage correction described in Section III reduces that remaining influence.

## III. Measuring Charge and Permittivity with Two Gates

### A. Chosen Dry-Readout Workflow

The proposed experimental workflow follows Kim *et al*.: immobilize the target, rinse away unbound material, dry the device and then record its electrical response [11]. Their biotin–streptavidin experiment used a deionized-water wash followed by nitrogen blow-drying before measurement. We adopt this dry-readout sequence; modeled *k* and interfacial charge describe the dry state.

### B. Stored Charge and Permittivity

Before measuring unknown samples, two reference curves are deliberately constructed using standards with known properties. For the BG reference, any biomolecule can in principle serve as the calibration material because the BG current depends only weakly on permittivity. Its current at 20 V is recorded for different known interfacial charges while keeping permittivity fixed; here, we use *k* = 5. For the TG reference, current at 7 V is recorded for neutral analytes with different known permittivities, where neutral means zero added interfacial charge.

The stored curves then translate BG current into charge and corrected TG current into permittivity by interpolation between neighboring calibration points. In this study, both curves are constructed from simulated currents at known charge and permittivity values, using the corresponding master-grid cases:

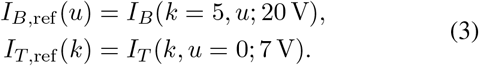

Here *u* = *N*_bio_*/*(10^12^ cm^*−*2^); all reported charge sweeps and plot charge coordinates refer to the added analyte contribution *N*_bio_. We denote the current-to-property lookup operations by *G*_*B*_ and *G*_*T*_, respectively. Both reference inversions use straight segments between sampled currents, extended by their nearest endpoint segment when necessary.

### C. First Reading: Estimate Charge

First, we measure the drain current with the bottom gate at 20 V and use the bottom-gate reference to estimate charge:

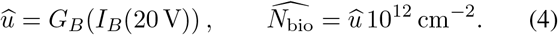

### D. Second Reading: Select the Corrected Top-Gate Voltage

Second, we measure the top-gate current at 4.5 V. This pilot current, *J*_4.5_ = |*I*_*T*_ (4.5 V)|*/*(1 *µ*A), and the estimated charge determine a top-gate voltage that compensates the remaining charge response:

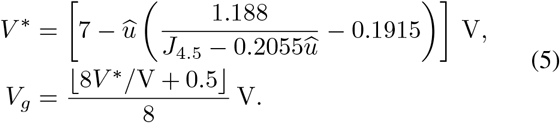

The pilot current selects the measurement voltage. The estimated charge and pilot current thus provide feedforward biasing, setting the top-gate voltage before the final permittivity measurement. The 7-V term is the stored neutral-reference bias. Because the simulated TG sweep uses 0.125-V increments, the calculated correction voltage is rounded to the nearest sampled bias, ensuring that extraction uses directly simulated currents. For zero estimated charge, the selected voltage is 7 V.

### E. Third Reading: Estimate Permittivity

Third, we measure the top-gate current at the selected voltage *V*_*g*_ and use the neutral reference to estimate permittivity:

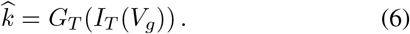

Voltage correction brings the third measured current close to the stored neutral-charge reference current for the same permittivity. We therefore interpret sample current measured at *V*_*g*_ using the original 7-V reference. These three readings estimate charge from the bottom gate and permittivity from the corrected top gate.

## IV. Simulation Model and Test Design

### A. TCAD Setup and Experimental Calibration

The proposed sensor is based on Zhang *et al*.’s asymmetric dual-gate oxide-TFT architecture [17], modified here by shortening the channel and incorporating two sensing nanogaps. We calibrated TCAD against their experimental transfer curves. The overlays in Fig. 2 reproduce the experimental responses for different bottom-gate-insulator thicknesses, drain biases and channel lengths. Agreement across these conditions supports the robustness of the calibrated model and its reproduction of the underlying transistor response. The calibrated model evaluates simultaneous charge and permittivity variation. ATLAS drift–diffusion simulations use Shockley–Read– Hall (SRH) recombination, bulk tail/Gaussian density-of-states (DOS) defects, additional 5-nm interface-adjacent defect slabs and Gummel–Newton trap handling. The localized volumetric defects are distinct from fixed interface sheet charge. The mesh was refined until the simulated drain-current values converged.

**Fig 2.**
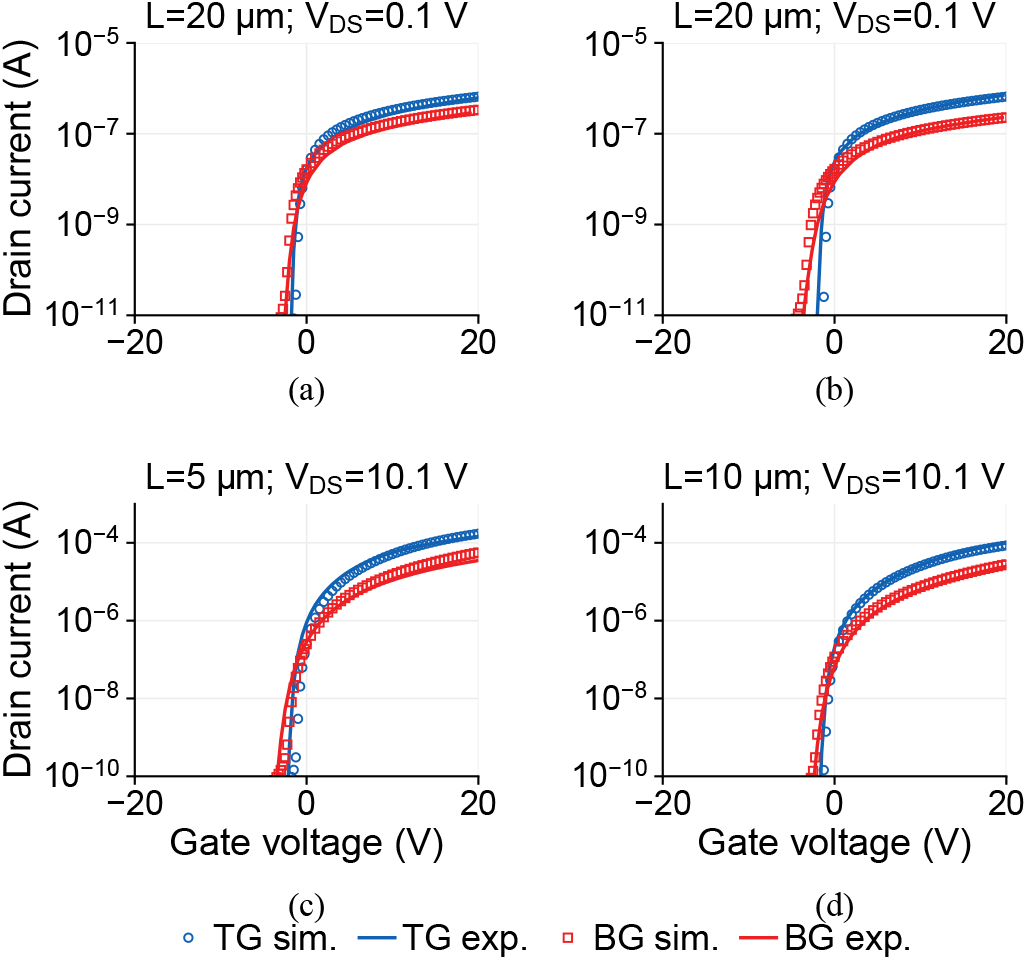
TCAD calibration against Zhang et al. [17]: (a,b) *L* = 20 *µ*m, bottom insulators of 200 and 300 nm, respectively, at *V*_*DS*_ = 0.1 V; (c,d) *L* = 5 and 10 *µ*m at *V*_*DS*_ = 10.1 V. Lines: experiment; markers: simulation.

**TABLE I.** Nominal model parameters.

| Parameter | Value |
| --- | --- |
| Top/bottom stacks | 20/500 nm |
| Nanogap thickness/length | 20 nm / 0.50 $\mu\text{m}$, each |
| HfO<sub>2</sub>/SiO<sub>2</sub> permittivity | 22 / 3.9 |
| $V_{DS}$; opposite gate | 0.1 V; 0 V |
| $T$; all electrode work functions | 300 K; 4.3 eV |
| Channel doping; $\mu_n/\mu_p$ | $10^{15} \text{ cm}^{-3}$; 9/$10^{-8} \text{ cm}^2/\text{Vs}$ |
| S/D doping; Gaussian length | $5 \times 10^{20} \text{ cm}^{-3}$; 0.015 $\mu\text{m}$ |
| Bulk $N_{TA}/N_{TD}$; $N_{GA}/N_{GD}$ | $5 \times 10^{18}/10^{19}$; $10^{16}/5 \times 10^{16}$ |
| Bulk $W_{TA}/W_{TD}$; $W_{GA}/W_{GD}$ | 0.015/0.12; 0.1/0.1 eV |
| Bulk $E_{GA}/E_{GD}$ | 0.4/0.4 eV |
| Local $N_{TA}/N_{TD}$; slab thickness | $10^{11}/5 \times 10^{11}$; 5 nm |
| Bottom/top baseline sheet charge | $4.60/4.80 \times 10^{11} \text{ cm}^{-2}$ |
| Solver; trap controls | Gummel-Newton; 0.5 / 8 |

### B. Development Grid and Analyte Loading Range

We simulated two datasets with the same geometry, material parameters, mesh, transport models and defect inputs. For every charge–permittivity pair, a bottom-gate sweep and a top-gate sweep were performed separately, with the opposite gate grounded and *V*_*DS*_ = 0.1 V.

The *master sweep* is the development dataset: every combination of ten permittivities and nine charges, giving 10 × 9 = 90 cases. It supplies the reference curves and the fitted voltage-correction coefficients. The grid is

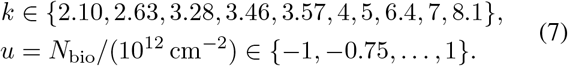

The normalized charge is *u* = *N*_bio_*/*(10^12^ cm^*−*2^); its nine values are −1, −0.75, −0.50, −0.25, 0, 0.25, 0.50, 0.75, 1.

Thus both properties vary together, including neutral, positively charged and negatively charged cases.

The full grid deliberately samples the entire intended operating domain, so its use to construct the references, fit a common correction law and assess full-range performance is part of the calibration design. Master-grid errors characterize this domain; coefficients remain fixed for off-grid tests.

The TCAD study covers 2.10 ≤ *k* ≤ 8.10 and −10^12^ ≤ *N*_bio_ ≤ 10^12^ cm^*−*2^, including zero charge. The ten effective dielectric loadings represent streptavidin (*k* = 2.10), biotin (2.63), cholesterol oxidase (3.28) and zein (5.00) [8], [18]; glucose oxidase (3.46) [19]; APTES (3.57), SARS-CoV-2 spike-protein loading (4.00) and gluten (7.00) [10]; and bacteriophage T7 (6.40) and apomyoglobin (8.10) [20].

We vary *k* and *N*_bio_ independently to evaluate charge– permittivity interference across a common electrical loading range. The nine *N*_bio_ levels describe effective interfacial sheet-charge inputs, whose experimental values depend on analyte chemistry, surface coverage and preparation conditions. Positive and negative sheet-charge densities on the 10^11^–10^12^ cm^*−*2^ scale are established modeling choices [8], [9]; Chahardah Cherik and Mohammadi explicitly examine ±5 × 10^11^ and ±10^12^ cm^*−*2^ [21]. The endpoints therefore sample the more strongly charged conditions within this literature-based range, alongside the smaller-magnitude and neutral states.

### C. Independent Holdout Test Design

The *holdout set* is a separate test dataset of 18 new charge– permittivity combinations. Both coordinates of every case were deliberately chosen between master-grid values, so none overlaps a calibration point. Reference curves and correction coefficients remained unchanged; no holdout case was used for fitting. This tests readout performance at unseen combinations beyond its calibration points. The (*k, u*) coordinates are (2.365, −0.920), (2.955, +0.680), (3.515, −0.420), (3.785, +0.120), (4.500, −0.120), (5.700, +0.420), (6.700, −0.680), (7.550, +0.920), (4.250, −0.800), (7.800, −0.550), (5.350, −0.300), (6.850, −0.180), (2.750, −0.060), (7.850, +0.060), (5.850, +0.180), (6.050, +0.300), (2.400, +0.550), (3.800, +0.800).

## V. Results: Gate Responses and Two-Property Extraction

### A. Bottom-Gate Charge Readout and Top-Gate Permittivity Response

Fig. 3(a) plots bottom-gate current at 20 V against charge for all ten permittivities. The curves nearly overlap on the full current scale: changing permittivity at fixed charge produces a maximum spread of only 0.0121 *µ*A. Consequently, a current read from the bottom-gate sweep can estimate charge without knowing the unknown sample’s permittivity.

**Fig 3.**
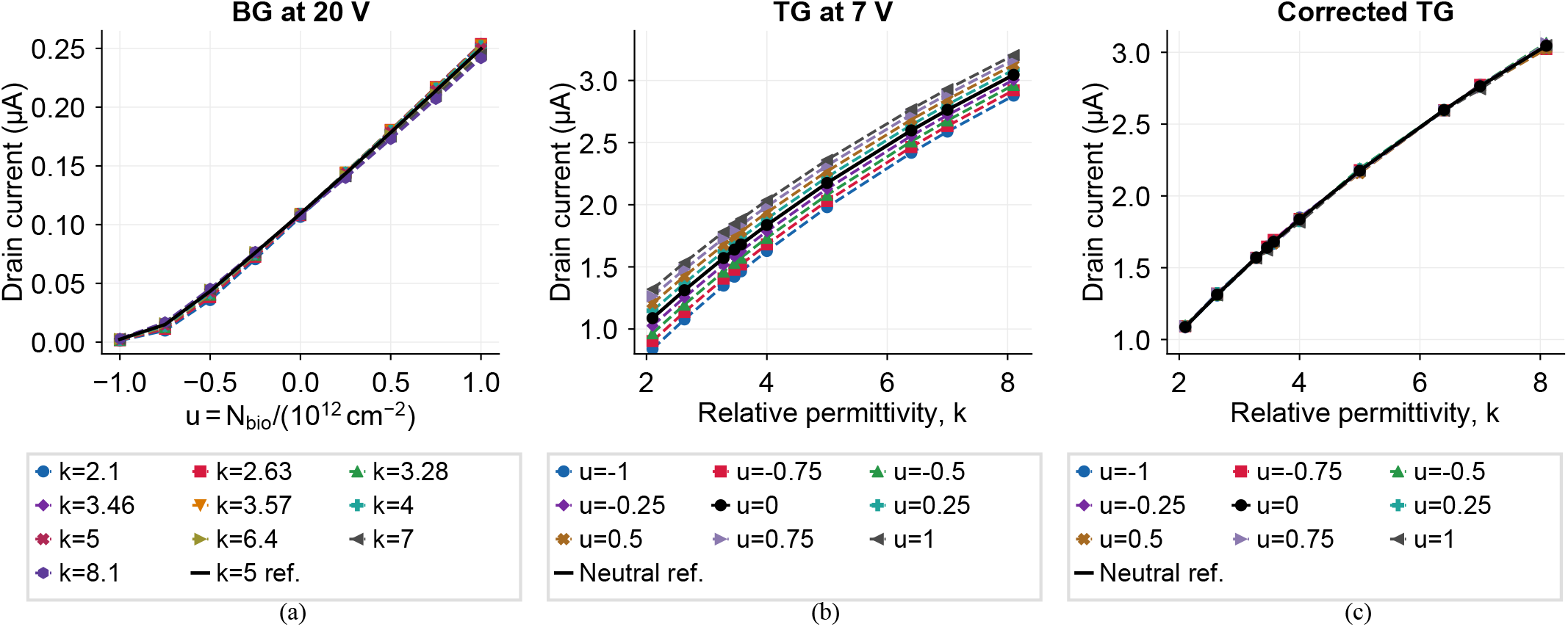
Master-grid current responses: (a) BG at 20 V, (b) TG at 7 V, and (c) corrected TG using (5). Black curves: *k* = 5 BG reference and unchanged neutral-charge TG reference at 7 V.

This close agreement allows one known-permittivity curve to serve as the charge reference. We use *k* = 5, near the midpoint 5.1 of the studied 2.1–8.1 range. It represents a central calibration standard.

Fig. 3(b) shows top-gate current at 7 V against permittivity for all nine charge levels. These responses are approximately linear and closely grouped, although they do not overlap completely. At each permittivity, the charge-induced current spread is 0.326–0.481 *µ*A. The top-gate sweep therefore provides the second property-sensitive reading, but residual charge dependence limits permittivity accuracy until the readout voltage is corrected.

### B. Improving the Top-Gate Permittivity Readout

Equation (5) uses the bottom-gate charge estimate and the 4.5-V pilot current to shift the top-gate readout around its 7-V reference bias. Fig. 3(c) shows the result: the charge-dependent curves closely collapse onto the unchanged neutral reference. Their spread falls to 0.0053–0.0424 *µ*A, or 0.20–1.93% of the neutral-reference current at the same permittivity. This close overlap allows the corrected current to estimate permittivity with little residual dependence on charge, completing the two-property readout. The optimized dielectric layers and half-channel nanogaps create the preferential gate responses; the correction provides further cross-sensitivity mitigation in the top-gate estimate. These overlays confirm accurate *k* recovery from one reference across charge states.

### C. Cross-Sensitivity Between the Two Property Measurements

The overlays demonstrate cross-sensitivity mitigation through the complementary gate responses. Fig. 4 quantifies the remaining cross-sensitivity across the full charge– permittivity range: a smaller ratio means that the desired property produces a stronger response than the interfering property. Each cell compares the response to the unwanted property with the response to the property being measured, after accounting for the studied parameter ranges:

**Fig 4.**
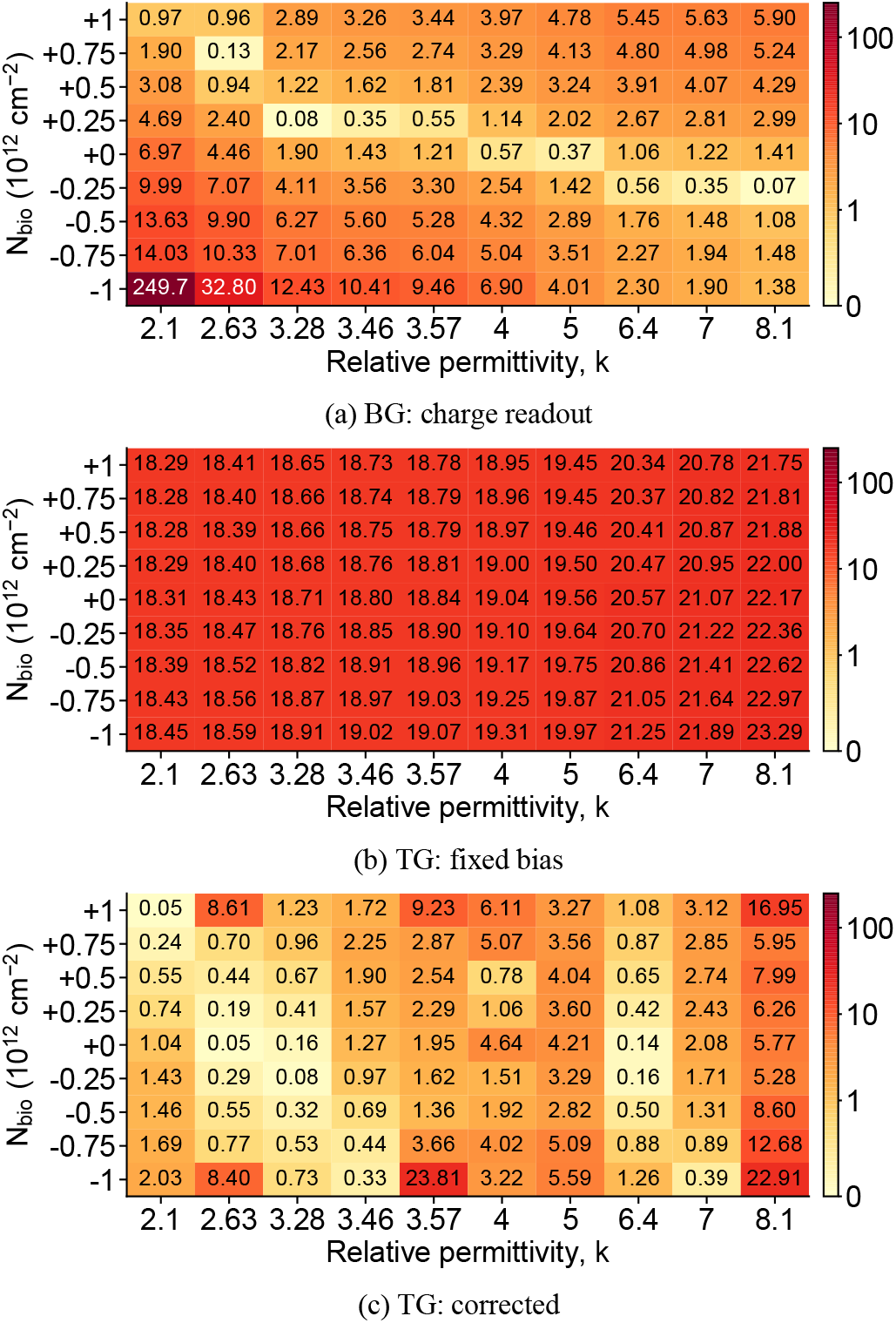
Span-normalized cross-sensitivity across 90 cases: (a) BG at 20 V, (b) TG at 7 V, and (c) corrected TG. The scale is linear below 1% and logarithmic above. Derivatives use nonuniform *k* spacing; panel (c) includes charge estimation and voltage rounding.

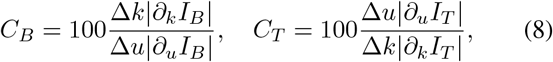

where Δ*k* = 6 and Δ*u* = 2. Smaller ratios mean stronger cross-sensitivity mitigation. Derivatives use second-order finite differences with the actual, nonuniform *k* spacing.

The bottom-gate mean/median ratios of 6.78%/3.04% show that charge dominates its response despite simultaneous permittivity changes. For the top gate, voltage correction reduces the mean/median ratios from 19.62%/18.98% to 3.05%/1.60%, substantially strengthening the permittivity readout. The device therefore supplies complementary responses before correction, and voltage selection further improves the second measurement.

A separate derivative check confirms that complementary responses exist before correction. Normalizing the inputs by their ranges and each fixed-bias current by its grid span gives

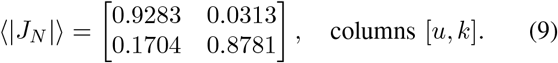

Rows represent bottom- and top-gate currents. Dominant diagonal entries confirm that the bottom gate predominantly responds to charge and the top gate to permittivity.

### D. Top-Gate Sensitivity for Permittivity-Based Identification

Permittivity provides a dielectric descriptor for distinguishing analytes with different effective *k*. Fig. 5 shows the corrected top-gate response relative to neutral air (*k* = 1, zero added analyte charge). We express this response as 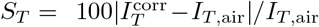, where *I*_*T*,air_ is the neutral-air current at 7 V. Across the studied cases, this air-referenced sensitivity ranges from 89.46% to 436.05%, demonstrating a large current contrast relative to air.

**Fig 5.**
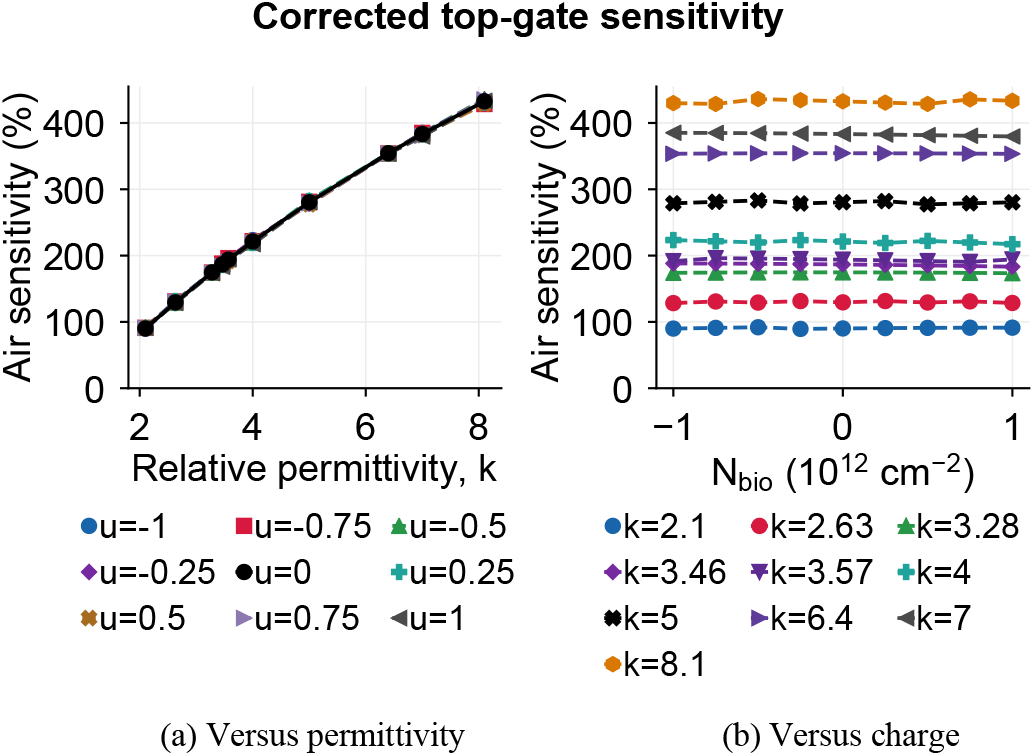
Corrected TG sensitivity relative to neutral air at 7 V: (a) versus permittivity and (b) versus charge, for all 90 cases.

**TABLE II.** Master-grid MEAN / RMS / worst errors (%), *N* = 90: absolute charge errors normalized to 10^12^ cm^*−*2^; absolute permit-tivity errors relative to true *k*.

| Readout | Mean | RMS | Worst |
| --- | --- | --- | --- |
| Charge | 1.743 | 2.438 | 10.183 |
| $k$, top gate at 7 V | 8.343 | 10.349 | 27.790 |
| $k$, corrected and rounded | 0.496 | 0.633 | 1.568 |

In panel (a), sensitivity increases approximately linearly with permittivity, and the curves for different charges nearly overlap. Panel (b) shows the same response remains nearly constant as charge varies at each permittivity. Thus the top gate retains a strong response to the dielectric property useful for analyte identification while substantially reducing charge interference. Alongside bottom-gate charge estimation, this yields two analyte descriptors.

### E. Accuracy on the 90-Case Master Grid

Charge and permittivity are estimated by matching measured currents to corresponding reference curves. For both datasets, the errors are

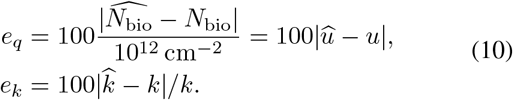

Fixed charge-error normalization includes neutral cases; permittivity error is relative to the true *k*. Table II reports the mean, root-mean-square (RMS) and largest errors across the 90 master cases. The grid was designed to cover the full operating range and supplies the references and fitted correction; these development-grid results therefore quantify full-range calibrated performance.

Mean charge error is 1.743%. The largest bottom-gate charge errors occur near the strongly negative boundary of the selected literature-based range. Smaller-magnitude charge cases yield smaller errors. Voltage correction lowers mean permittivity error from 8.343% at a fixed top-gate bias of 7 V to 0.496%, a 94.06% reduction. These results show that the calibrated device-and-readout strategy measures both properties across the deliberately sampled operating domain. The following section tests the unchanged method on independent combinations.

## VI. Independent Validation and Noise Assessment

### A. Accuracy on the 18 Independent Holdout Cases

The 18 holdouts defined in Section IV-C were deliberately placed between calibration points in both charge and permittivity. They are evaluated using the unchanged master-derived references and correction equation, without refitting. Fig. 6 compares their currents with those references, and Table III reports errors calculated using (10).

**TABLE III.**
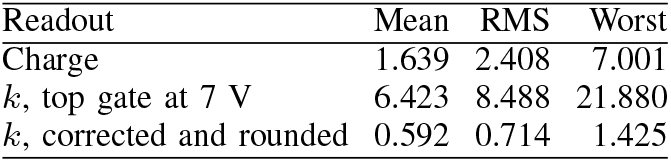
Independent holdout mean / RMS / worst errors (%), *N* = 18: absolute charge errors normalized to 10^12^ cm^*−*2^; absolute permittivity errors relative to true *k*.

| Readout | Mean | RMS | Worst |
| --- | --- | --- | --- |
| Charge | 1.639 | 2.408 | 7.001 |
| $k$, top gate at 7 V | 6.423 | 8.488 | 21.880 |
| $k$, corrected and rounded | 0.592 | 0.714 | 1.425 |

**Fig 6.**
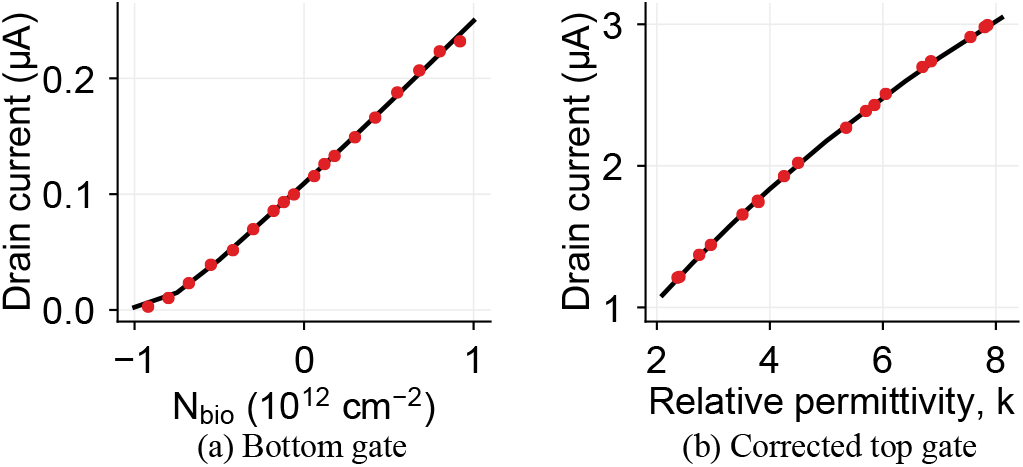
Independent holdout currents: (a) BG at 20 V and (b) corrected TG. Black curves: frozen *k* = 5 BG and neutral-charge TG references; red circles: 18 holdouts. The TG reference is at 7 V.

The mean charge error is 1.639%. As in the master grid, the largest bottom-gate deviations occur toward the strongly negative end of the tested charge range; the interior charge cases provide better extraction accuracy. Mean relative permittivity error falls from 6.423% at a fixed 7-V top-gate bias to 0.592% after correction; the largest corrected error is 1.425%. Thus the method retains accurate charge and permittivity extraction at combinations not used for calibration or fitting.

### B. Supporting Intrinsic-Noise Check

ATLAS noise analysis covered 12 charge–permittivity pairs (*k* ∈ {2.10, 3.57, 5, 8.10}, *u* ∈ {−1, 0, +1}) at a fixed bottom-gate bias of 20 V and a fixed top-gate bias of 7 V, respectively, with the opposite gate grounded and *V*_*DS*_ = 0.1 V. Trapezoidal integration of the total drain-current PSD over 1 Hz–100 kHz gave RMS noise *σ*_*I*_ of 0.029–0.211 nA for the bottom gate and 0.639–0.837 nA for the top gate. For independent readings, the sampled separation ratio 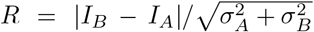 had minima of 539.9 and 412.5, respectively (Fig. 7). Thus, modeled intrinsic noise is negligible relative to the sampled current separations.

**Fig 7.**
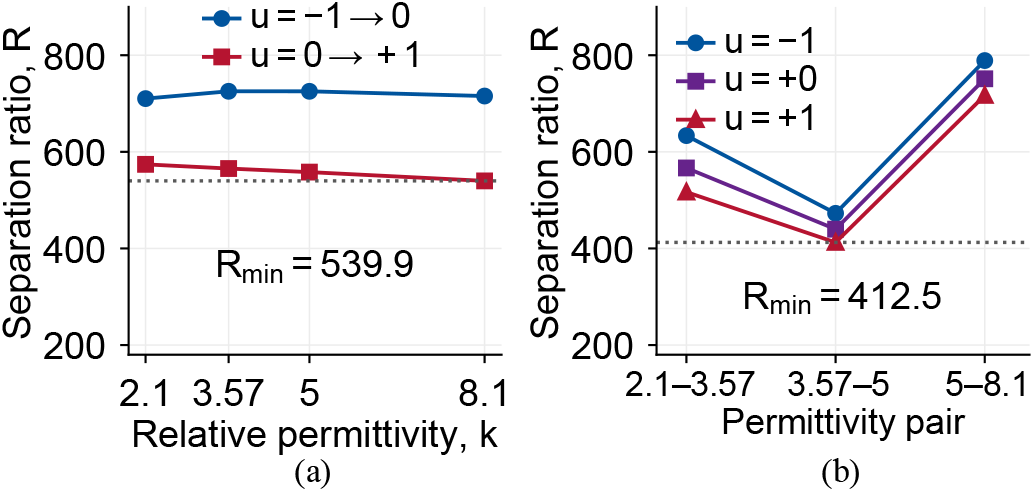
Current separation relative to combined RMS noise over 1 Hz–100 kHz: (a) adjacent charges at fixed *k*, BG at 20 V; (b) adjacent permittivities at fixed *u*, TG at 7 V. Dotted lines mark minimum ratios.

## VII. Conclusion

This conceptual demonstration uses the two gates of one asymmetric dual-gate, dual-nanogap a-IGZO transistor to estimate two effective analyte properties: charge from the bottom-gate current reading and permittivity from the top-gate current reading. Both properties vary simultaneously in both sensing regions. The unequal dielectric stacks and optimized half-channel nanogaps give the gate currents their preferential measurement roles; charge-based selection of the top-gate voltage improves the second estimate. Using the bottom-gate reference at 20 V and *k* = 5 and the neutral-charge top-gate reference at 7 V gives mean charge errors of 1.74% and 1.64% for the master and holdout sets, with corresponding permittivity errors of 0.50% and 0.59%. All nominal corrected top-gate readings are directly logged. Device-level cross-sensitivity mitigation and voltage correction address the ambiguity of assigning a charge-dependent current change to permittivity alone. This enables accurate permittivity measurement across positive, negative and zero analyte charge within the tested range, alongside simultaneous two-property estimation and physics-based dielectric-layer and nanogap engineering. The resulting contribution is a novel device-and-readout strategy for measuring both properties from coupled charge and dielectric responses through sequential current measurements, establishing a numerical basis for future biosensing studies.

## Acknowledgment

ChatGPT (OpenAI) was used to assist with manuscript and figure preparation, numerical post-processing, and the regeneration of TCAD scripts for the extensive simulation matrix.

## References

[1] H. Im, X.-J. Huang, B. Gu, and Y.-K. Choi, “A dielectric-modulated field-effect transistor for biosensing,” Nat. Nanotechnol., vol. 2, no. 7, pp. 430–434, 2007, doi: 10.1038/nnano.2007.180.

[2] M. Im, J.-H. Ahn, J.-W. Han, T. J. Park, S. Y. Lee, and Y.-K. Choi, “Development of a point-of-care testing platform with a nanogap-embedded separated double-gate field effect transistor array and its readout system for detection of avian influenza,” IEEE Sensors J., vol. 11, no. 2, pp. 351–360, Feb. 2011, doi: 10.1109/JSEN.2010.2062502.

[3] E. Stern, R. Wagner, F. J. Sigworth, R. Breaker, T. M. Fahmy, and M. A. Reed, “Importance of the Debye screening length on nanowire field effect transistor sensors,” Nano Lett., vol. 7, no. 11, pp. 3405–3409, 2007, doi: 10.1021/nl071792z.

[4] S. Santermans et al., “The significance of nonlinear screening and the pH interference mechanism in field-effect transistor molecular sensors,” ACS Sensors, vol. 6, no. 3, pp. 1049–1056, 2021, doi: 10.1021/acssensors.0c02285.

[5] R. Narang, K. V. S. Reddy, M. Saxena, R. S. Gupta, and M. Gupta, “A dielectric-modulated tunnel-FET-based biosensor for label-free detection: Analytical modeling study and sensitivity analysis,” IEEE Trans. Electron Devices, vol. 59, no. 10, pp. 2809–2817, Oct. 2012, doi: 10.1109/TED.2012.2208115.

[6] R. Narang, M. Saxena, and M. Gupta, “Comparative analysis of dielectric-modulated FET and TFET-based biosensor,” IEEE Trans. Nanotechnol., vol. 14, no. 3, pp. 427–435, May 2015, doi: 10.1109/TNANO.2015.2396899.

[7] D. Singh, B. S. Sengar, P. Dwivedi, and V. Garg, “Comparative analysis of gate structure dependent FET-based biosensor,” Mater. Today Commun., vol. 35, Art. no. 106301, Jun. 2023, doi: 10.1016/j.mtcomm.2023.106301.

[8] D. Srikanya, C. Sahu, G. Gopal, A. K. Johar, and T. Varma, “Dielectric modulated a-IGZO TFT biosensor: A cost-effective design for enhanced biomolecule detection,” Phys. Scr., vol. 100, no. 12, Art. no. 125021, 2025, doi: 10.1088/1402-4896/ae2bb6.

[9] D. Chowdhury et al., “A novel dielectric modulated gate-stack double-gate metal-oxide-semiconductor field-effect transistor-based sensor for detecting biomolecules,” Sensors, vol. 23, no. 6, Art. no. 2953, 2023, doi: 10.3390/s23062953.

[10] S. Khan and E. Rahman, “Designing the cavity architecture in double gate junctionless field effect transistors for enhanced biomolecule detection,” Nanoscale Adv., vol. 7, pp. 3746–3763, 2025, doi: 10.1039/D4NA00928B.

[11] S. Kim, D. Baek, J.-Y. Kim, S.-J. Choi, M.-L. Seol, and Y.-K. Choi, “A transistor-based biosensor for the extraction of physical properties from biomolecules,” Appl. Phys. Lett., vol. 101, no. 7, Art. no. 073703, 2012, doi: 10.1063/1.4745769.

[12] A. Cuervo, P. D. Dans, J. L. Carrascosa, M. Orozco, G. Gomila, and L. Fumagalli, “Direct measurement of the dielectric polarization properties of DNA,” Proc. Natl. Acad. Sci. USA, vol. 111, no. 35, pp. E3624– E3630, 2014, doi: 10.1073/pnas.1405702111.

[13] X. P. A. Gao, G. Zheng, and C. M. Lieber, “Subthreshold regime has the optimal sensitivity for nanowire FET biosensors,” Nano Lett., vol. 10, no. 2, pp. 547–552, 2010, doi: 10.1021/nl9034219.

[14] S. K. Das, S. M. Biswal, L. I. Giri, and D. Swain, “Thermal influence on performance characteristics of double gate MOSFET biosensors with gate stack configuration,” Discov. Appl. Sci., vol. 6, Art. no. 447, 2024, doi: 10.1007/s42452-024-06055-1.

[15] M. Kumari, N. K. Singh, and M. Sahoo, “A detailed investigation of dielectric-modulated dual-gate TMD FET based label-free biosensor via analytical modelling,” Sci. Rep., vol. 12, Art. no. 21115, 2022, doi: 10.1038/s41598-022-24677-6.

[16] N. Kannan and M. Jagadesh Kumar, “Dielectric-modulated impact-ionization MOS transistor as a label-free biosensor,” IEEE Electron Device Lett., vol. 34, no. 12, pp. 1575–1577, Dec. 2013, doi: 10.1109/LED.2013.2283858.

[17] Y. Zhang, L. Chang, H. Yang, L. Lu, C. Liao, and S. Zhang, “Asymmetric double-gate (ADG) oxide TFTs with both gradual and steep subthreshold characteristics for mobile AMOLED displays,” Displays, vol. 93, Art. no. 103453, 2026, doi: 10.1016/j.displa.2026.103453.

[18] P. Kumar and B. Raj, “Performance assessment and optimization of vertical nanowire TFET for biosensor application,” Trans. Electr. Electron. Mater., vol. 23, pp. 685–692, 2022, doi: 10.1007/s42341-022-00405-9.

[19] M. Patil, A. Gedam, and G. P. Mishra, “Performance assessment of a cavity on source charge-plasma TFET-based biosensor,” IEEE Sensors J., vol. 21, no. 3, pp. 2526–2532, Feb. 2021, doi: 10.1109/JSEN.2020.3027031.

[20] A. Theja and M. Panchore, “Performance investigation of GaSb/Si heterojunction based gate underlap and overlap vertical TFET biosensor,” IEEE Trans. Nanobiosci., vol. 22, no. 2, pp. 284–291, Apr. 2023, doi: 10.1109/TNB.2022.3183934.

[21] I. Chahardah Cherik and S. Mohammadi, “Fringe-fields-modulated double-gate tunnel-FET biosensor,” Sci. Rep., vol. 14, Art. no. 168, 2024, doi: 10.1038/s41598-023-50723-y.

